## supplemental figures for "Nkx2-5 defines distinct scaffold and recruitment phases during formation of the cardiac Purkinje fiber network"

Choquet *et al.*

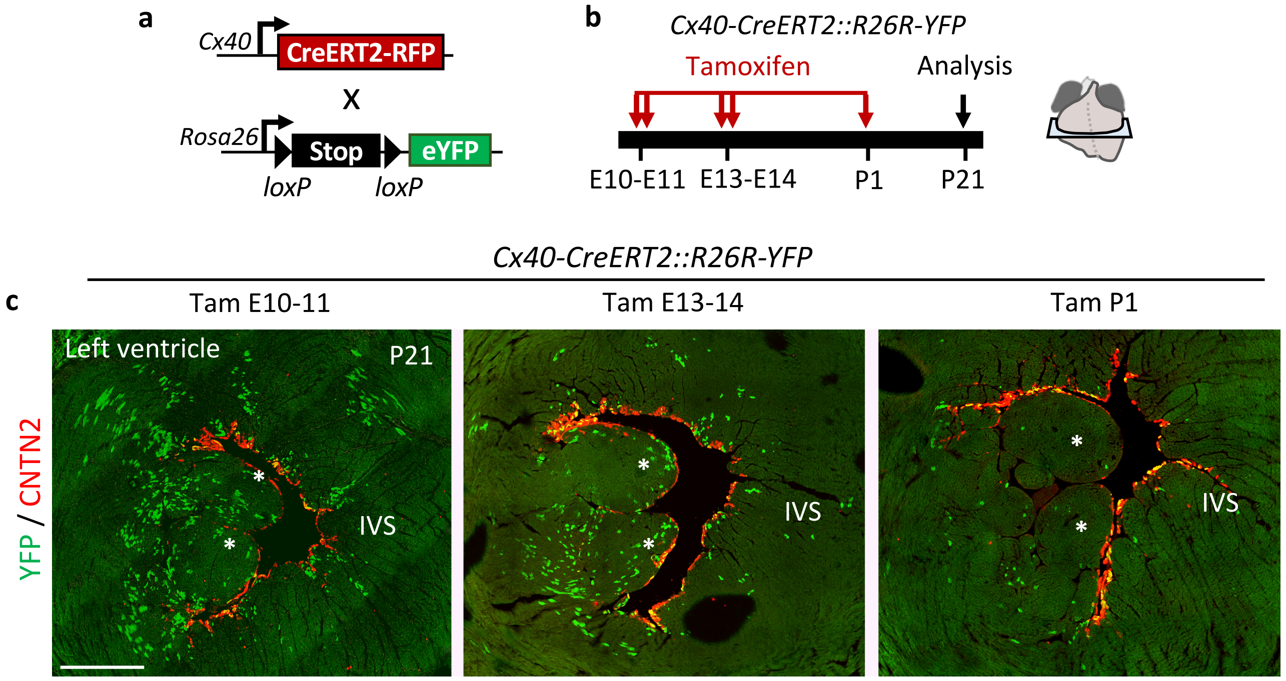


**Supplementary Figure 1**: **Inducible genetic tracing of *Cx40+* derivative cells during embryonic development or postnatal stages**. **a,** Schematic illustration of the genetic lineage tracing strategy. *R26R-YFP* reporter mouse line is crossed with tamoxifen inducible *Cx40-CreERT2* mice. Injection of tamoxifen induces the recombination of loxp sites resulting in the expression of the fluorescent protein YFP in *Cx40+*cells. **b**, Time course of tamoxifen injections and analyses of *Cx40+* derived-cells are represented on a time scale. Analysis are made on transverse sections in the middle region of ventricles. **c,** Immunostaining for YFP and CNTN2 on *Cx40-CreERT2::R26R-YFP* heart transverse sections at P21, show progressive restriction of *Cx40+* derived-cells (YFP+) to the VCS (CNTN2+) during ventricular morphogenesis. Asterisks represent papillary muscles; IVS: interventricular septum. Scale bar= 500µm. (n=4 Ctrl and *Nkx2-5*$^{+/-}$)

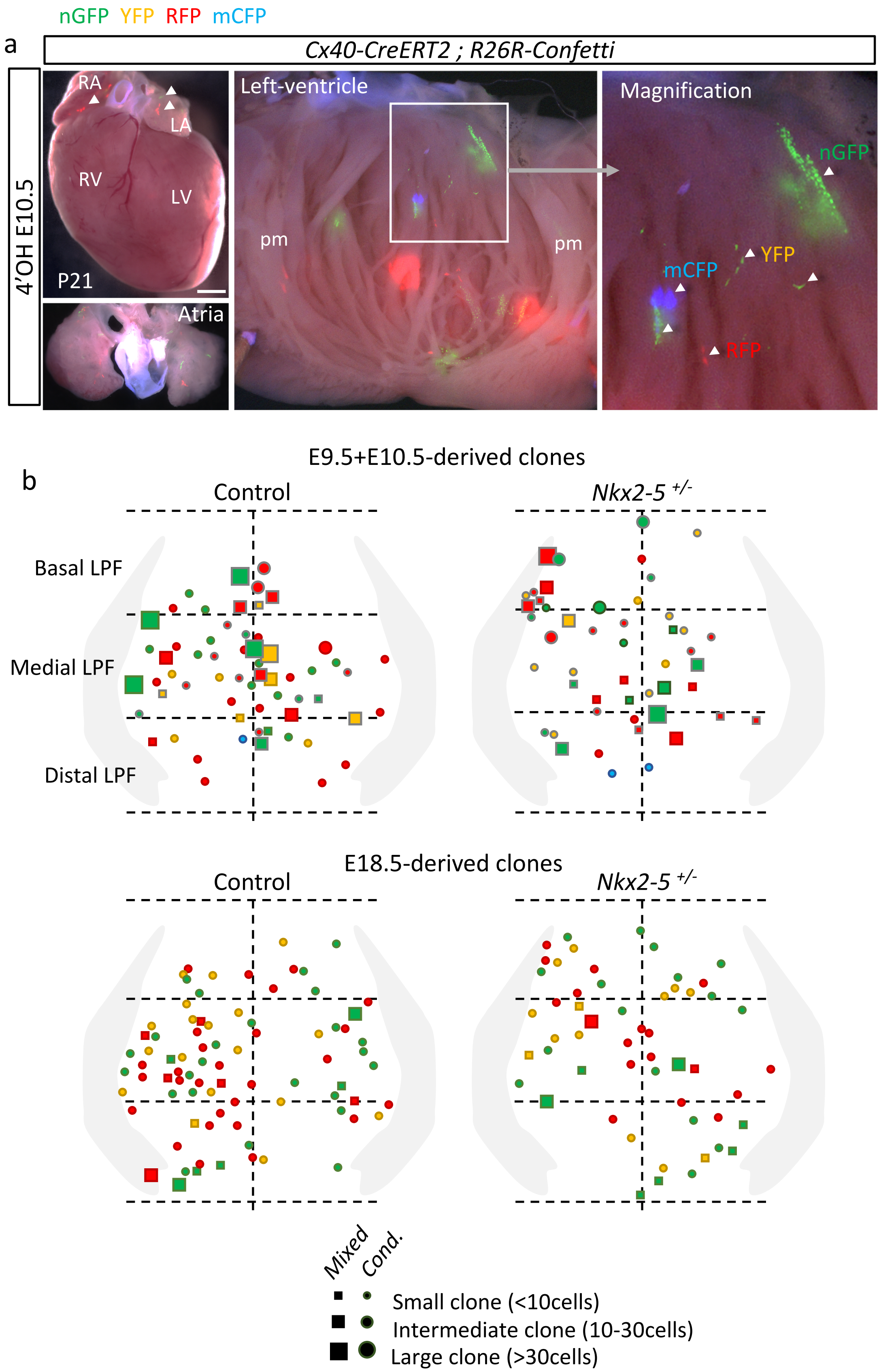


**Supplementary Figure 2: Whole-mount fluorescence views of P21 hearts with multicolor clusters from *R26R-Confetti* reporter line.**

**a,** Whole-mount views of an *Cx40-CreERT2*::*R26R-Confetti* heart induced by 4'OH injection at E10.5. The four fluorescent reporters (mCFP, YFP, nGFP, RFP) form independent unicolor clones distributed in the atria and along the left ventricular part of interventricular septum (IVS). Injection of 4'OH induces the recombination of loxp sites resulting in the expression of the fluorescent protein YFP in *Cx40+* cells. pm: papillary muscle; arrow heads show individual clones in atria and left ventricular myocardium. Scale bars = 1mm. (n=16 Ctrl; n=10 *Nkx2-5*$^{+/-}$) **b, Spatial location of clones within the IVS left-ventricular domain.** Schemas represent the spatial distribution of the total mixed (squares) and conductive (rounds) clones analyzed along the apico-basal axis. Each clone sizes are represented. The two first panels represent total E9.5 and E10.5-derived clones analyzed in control and *Nkx2-5*$^{+/-}$ hearts. The two lower panels represent total E18.5-derived clones analyzed in control and *Nkx2-5*$^{+/-}$ hearts.

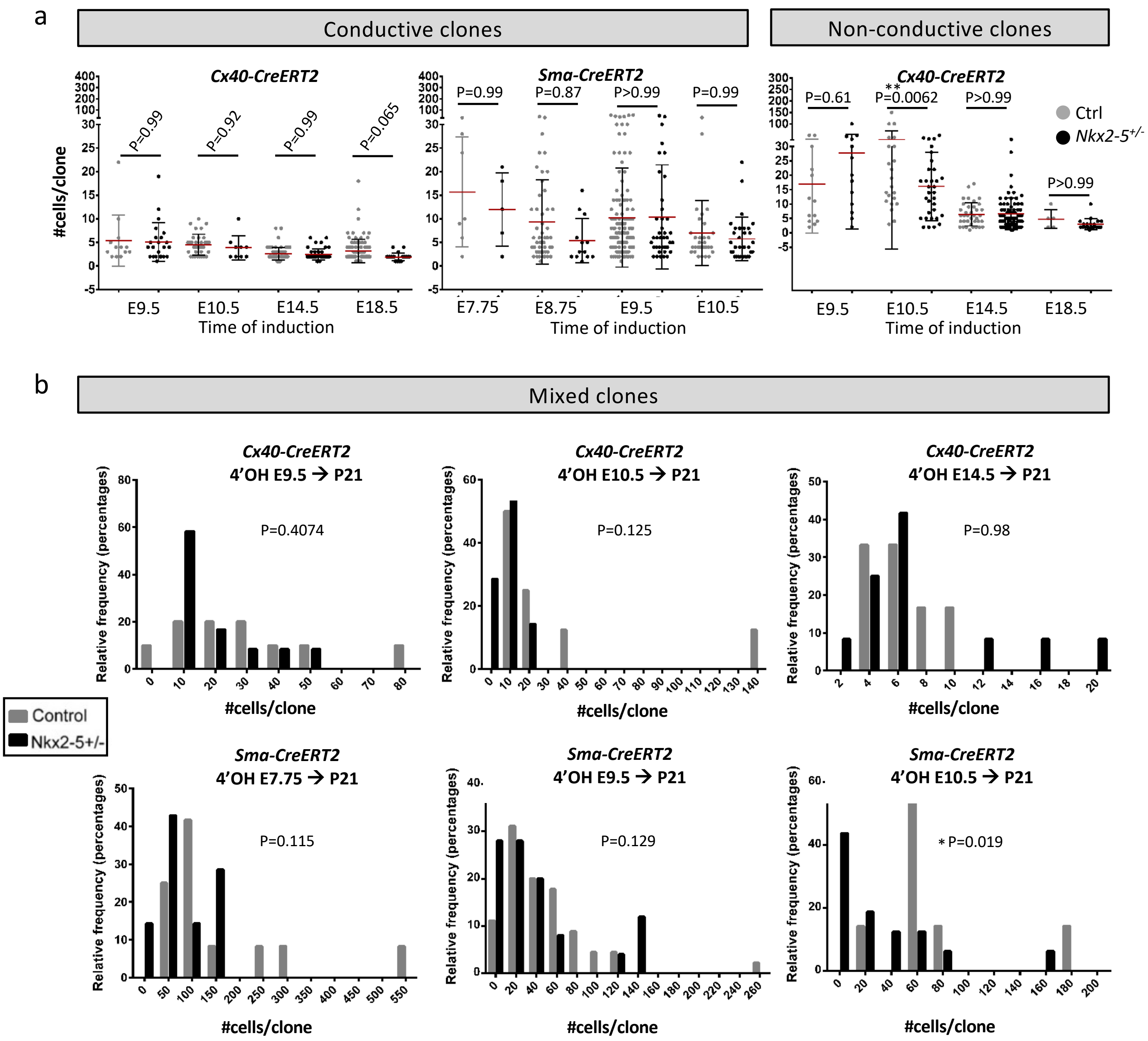


**Supplementary Figure 3: Size of temporally distinct *Cx40+* and *Sma+* derived-clones. a,** Dot plot graphs represent cell number per conductive and non-conductive individual clones. Quantification is made in P21 control (ctrl) or *Nkx2-5*$^{+/-}$ mice and after 4'OH induction using either *Cx40-CreERT2*::*R26R-Confetti* or *Sma-CreERT2*::*R26R-Confetti* mouse lines. Each dots represent individual clone. Red bars represent the Mean ± *SD*. *P* values are derived ordinary one-way ANOVA test (Tukey test),***P*<0.01. **b**, Graphs represent distribution of mixed clone size after induction at different time point. *P* value is derived from non-parametric Mann-Withney *t*-Test, **p<0.05*.

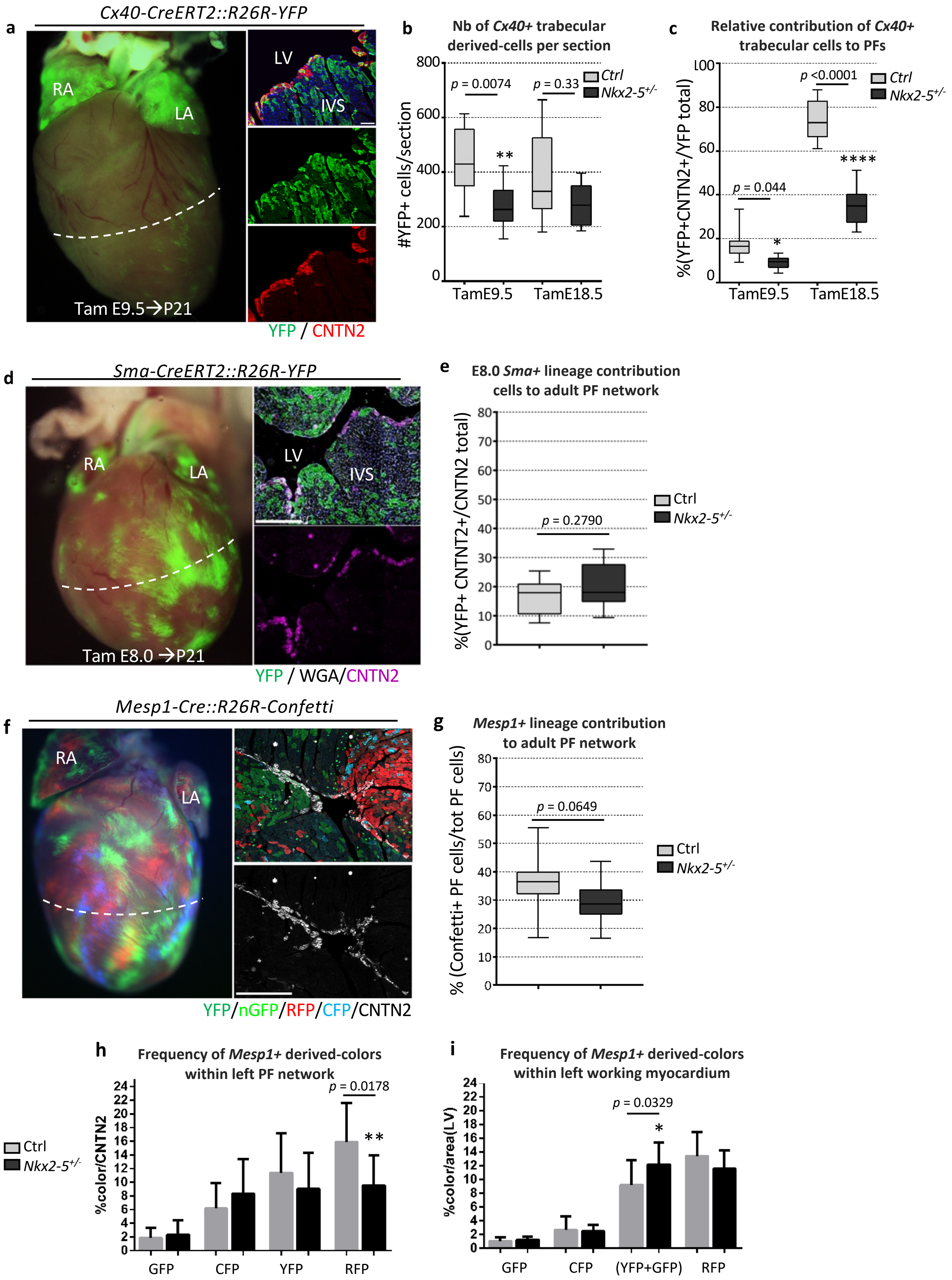
a
Cx40-CreERT2::R26R-YFP
RA
LA
LV
IVS
Tam E9.5→P21
YFP / CNTN2
b
Nb of Cx40+ trabecular derived-cells per section
#YFP+ cells/section
p = 0.0074
p = 0.33
**
Ctrl
Nkx2-5+/-
TamE9.5
TamE18.5
c
Relative contribution of Cx40+ trabecular cells to PFs
%(YFP+CNTN2+/YFP total)
p <0.0001
p = 0.044
*
****
Ctrl
Nkx2-5+/-
TamE9.5
TamE18.5
d
Sma-CreERT2::R26R-YFP
RA
LA
LV
IVS
Tam E8.0 →P21
YFP / WGA/CNTN2
e
E8.0 Sma+ lineage contribution cells to adult PF network
%(YFP+ CNTNT2+/CNTN2 total)
p = 0.2790
Ctrl
Nkx2-5+/-
f
Mesp1-Cre::R26R-Confetti
RA
LA
YFP/nGFP/RFP/CFP/CNTN2
g
Mesp1+ lineage contribution to adult PF network
% (Confetti+ PF cells/tot PF cells)
p = 0.0649
Ctrl
Nkx2-5+/-
h
Frequency of Mesp1+ derived-colors within left PF network
%color/CNTN2
p = 0.0178
**
Ctrl
Nkx2-5+/-
GFP
CFP
YFP
RFP
i
Frequency of Mesp1+ derived-colors within left working myocardium
%color/area(LV)
p = 0.0329
*
GFP
CFP
(YFP+GFP)
RFP

**Supplementary Figure 4**: **Contribution of different progenitors population to the VCS. a,** Whole-mount view of an P21 *Cx40-CreERT2*::*R26R-YFP* heart induced by tamoxifen injection at E9.5. Right panels show high magnification of left ventricular transverse sections after immunostaining using CNTN2 antibody. **b-c,** Graphs represent the quantification from (a) images. The total number of trabecular derived cells (YFP+) **(b)** and the relative contribution of trabecular cells to PFs ((YFP+CNTN2+)/YFP+) **(c)** are quantified in middle sections. Mean of 4 sections/heart TamE9.5 and 3sections/heart TamE18.5. (*n*=3 TamE9.5 Ctrl and TamE18.5 Ctrl and *Nkx2-5*$^{+/-}$; *n*=4 TamE9.5 *Nkx2-5*$^{+/-}$). *P* values are derived ordinary one-way ANOVA test,**P*<0.05; ***P*<0.01; *****P*<0.0001. Scale bar = 50µm. **d,** Whole-mount view of an P21 *Sma-CreERT2::R26R-YFP* heart induced by tamoxifen injection at E8.0. Right panels show high magnification of left ventricular transverse sections after immunostaining using CNTN2 and WGA antibodies. **e,** Graph represents the percentage of YFP-labelled adult PFs according to *Sma+* lineage traced by Tam injection at E8.0( YFP+CNTN2+)/CNTN2+). Quantifications are made from (d) images using control (ctrl) or *Nkx2-5*$^{+/-}$ heart transverse sections. Mean of 3sections/heart; (*n*=4 Ctrl; *n*=3 *Nkx2-5*$^{+/-}$). *P* values are derived ordinary one-way ANOVA test (Tukey test). **f,** Whole-mount view of an P21 *Mesp1-CreERT2::R26R-Confetti*. Right panels show high magnification of left ventricular transverse sections after immunostaining using CNTN2 antibody. **g,** Graph represents the percentage of Confetti-labelled adult PFs according to *Mesp1+* lineage (Confetti+CNTN2+)/CNTN2+). Quantifications are made from (f) images using control (ctrl) or *Nkx2-5*$^{+/-}$ heart transverse sections. Mean of 3sections/heart; (*n*=4 Ctrl and *Nkx2-5*$^{+/-}$). *P* values are derived ordinary one-way ANOVA test (Tukey test). **h-i,** Diagrams represent the individual *Mesp1+ c*onfetti colors percentage contribution to the left Purkinje network (h) or the working myocardium (i) in control or *Nkx2-5*$^{+/-}$ hearts (*n*=4). *P* values are derived ordinary one-way ANOVA test (Tukey test),**P*<0.05; ***P*<0.01.

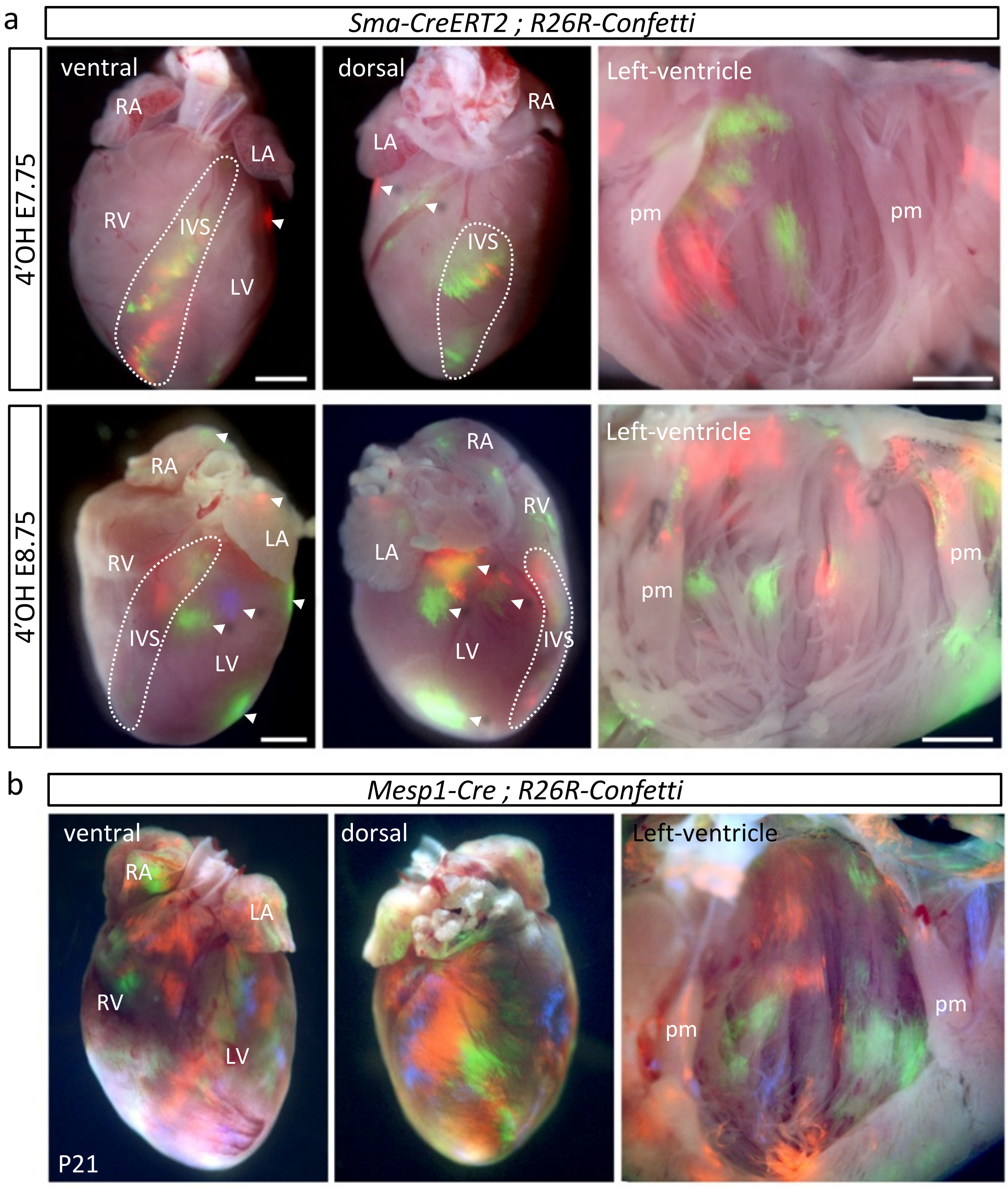


**Supplementary Figure 5: Whole-mount fluorescence views of P21 hearts with multicolor clusters from *R26R-Confetti* reporter line. a,** Whole-mount views of *Sma-CreERT2*::*R26R-Confetti* hearts injected at E7.75 or E8.75. The E7.75 *Sma+* derived clones are mostly distributed within the IVS and some in the left ventricle. The E8.75 *Sma+* derived clones are distributed within the IVS, the left ventricle, the atria and some within the right ventricle. **b,** Whole-mount views of *Mesp1-Cre*::*R26R-Confetti* hearts. Clones derived form *Mesp1* expression (E6.25 to E7.5) participate to the different regions of the heart. Scale bars= 1mm.

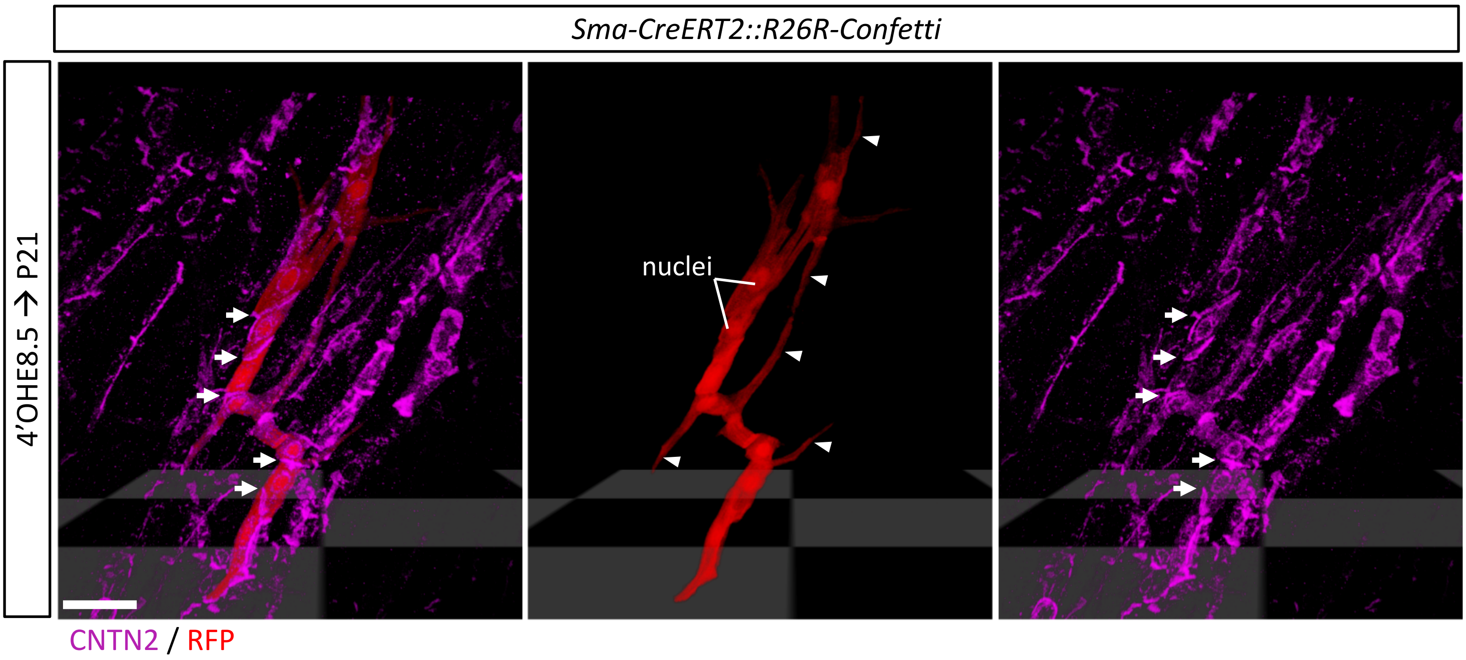


**Supplementary Figure 6: 3D confocal imaging reconstruction of a conductive clone induced during embryonic stages and observed in P21 left Purkinje fiber network**. RFP+ clone from *R26R-Confetti::Sma-CreERT2* crossing after 4'OH injection at E8.5 helps to appreciate the cellular morphology of conductive cells that present several cell extensions (arrow heads) within the left ventricular PF network. Whole-mount immunostaining for CNTN2 shows accumulation of CNTN2 within peripheral membrane of the conductive cell body that establish contact with other conductive cells (arrows). CNTN2 is less present within the cell extensions of conductive cells. Scale bar = 50µm

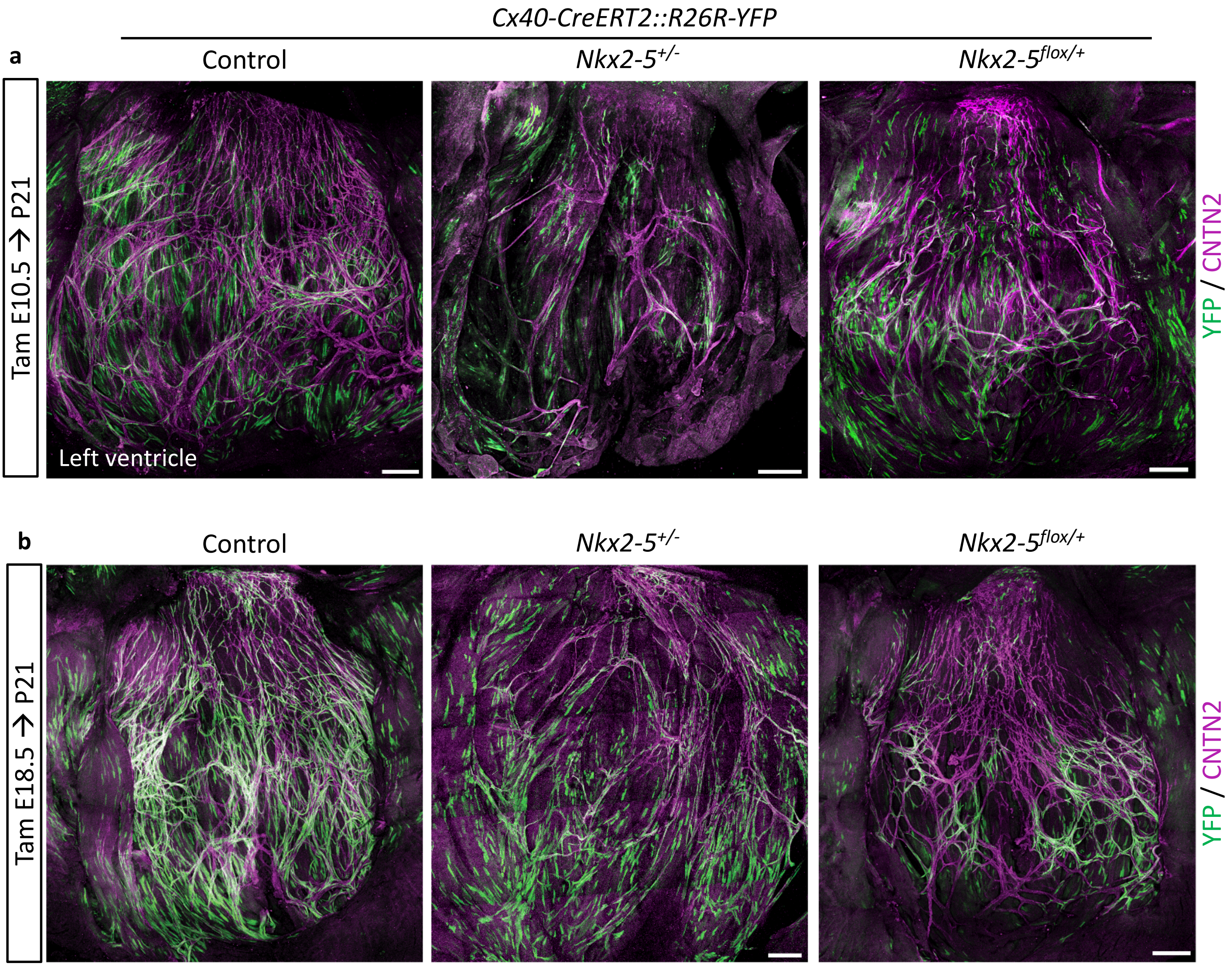


**Supplementary Figure 7: *Cx40+* lineage tracing reveals that a maximal level of Nkx2-5 is required during embryonic stages for proper Purkinje network development. a-b,** Confocal imaging of whole-mount immunostaining of *Cx40-CreERT2::R26R-YFP* opened-left ventricles at P21, after E10.5 (a) or E18.5 (b) tamoxifen injection. Immunostaining for YFP and CNTN2 on control, *Nkx2-5*$^{+/-}$ or *Nkx2-5*$^{flox/+}$ mice show affected participation of *Cx40+* derived-cells to the PF network when Nkx2-5 expression is reduced during E10.5 embryonic stage. Scale bars=500µm
